# High resolution imaging and interpretation of three-dimensional RPE sheet structure

**DOI:** 10.1101/2024.12.04.626881

**Authors:** Kevin J. Donaldson, Micah A. Chrenek, Jeffrey H. Boatright, John M. Nickerson

## Abstract

The retinal pigment epithelium (RPE), a monolayer of pigmented cells, is critical for visual function through its interaction with the neural retina. In healthy eyes, RPE cells exhibit a uniform hexagonal arrangement, but under stress or disease, such as age-related macular degeneration (AMD), dysmorphic traits like cell enlargement and apparent multinucleation emerge. Multinucleation has been hypothesized to result from cellular fusion, a compensatory mechanism to maintain cell to cell contact, barrier function, and conserve resources in unhealthy tissue. However, traditional two-dimensional (2D) imaging using apical border markers alone may misrepresent multinucleation due to the lack of lateral markers.

We present high-resolution confocal images enabling three-dimensional (3D) visualization of apical (ZO-1) and lateral (alpha-catenin) markers alongside nuclei. In two RPE damage models, we find that seemingly multinucleated cells are often single cells with displaced neighboring nuclei and lateral membranes. This emphasizes the need for 3D analyses to avoid misinterpreting multinucleation and underlying fusion mechanisms. Lastly, images from the NaIO3 oxidative damage model reveal variability in RPE damage, with elongated, dysmorphic cells showing increased ZsGreen reporter protein expression driven by EMT-linked CAG promoter activity, while more regular RPE cells displayed somewhat reduced green signal more typical of epithelial phenotypes.

## Introduction

The retinal pigment epithelium (RPE) is a monolayer sheet of cells in the eye that forms a barrier between the neural retina and the choroid. The RPE plays a critical role in retinal homeostasis through a myriad of functions, primarily physically and metabolically supporting the adjacent rod and cone photoreceptor cells of the neural retina (Bhatia et al., 2016a; Rashid et al., 2016; Strauss, 2005; Wang et al., 2018; Q. Zhang et al., 2019).

In healthy RPE tissue, the apical surfaces of individual cells in the sheet present with the appearance of regular hexagons when viewed *en face*. Each RPE cell typically makes contact with six immediately adjacent neighbor RPE cells, establishing the outer blood retina barrier. RPE cells are terminally differentiated and do not divide in either adult human subjects or rodent model species.

Studies examining RPE cell structure have revealed correlations between abnormal morphology and multiple aging and blindness disorders, such as age-related macular degeneration (AMD) and other inherited retinopathies (Marmor & Wolfensberger, 1998; Gambril et al., 2019; Bonilha, 2008; Christensen et al., 2017; Michaelides et al., 2003). Under stressful conditions, the RPE sheet, which is generally highly robust, can undergo numerous signs of damage, exhibiting both functional and morphological changes. While the RPE sheet can lose barrier functions, it appears to preserve the monolayer barrier by employing multiple reconfiguration processes. For instance, dead or dying RPE cells can be extruded in a classical purse-string mechanism, and surrounding cells appear to fill in by changing shapes in a pie-like pattern (Hergott et al., 1989; Kalnins et al., 1995).

Under aging conditions, as well as after severe stress, some RPE cells become much larger, but it is unclear how this happens. Is it a single cell that gains size, or possibly two cells fused together, resulting in a larger single cell? Additionally, it is unclear how these stressed or damaged cells are able to move. Is there a modification to Bruch’s membrane itself, such that cells translocate by traversing laterally on the surface of a stationary Bruch’s membrane, or is it the reverse, where perhaps Bruch’s membrane is modified itself and the attached anchored cells move as the membrane does?

An additional observation of these cells under abnormal conditions is that some appear to be multinucleate. While it is well established that rodent RPE are uni- or binucleate, it is less clear if, or how, a single RPE cell might acquire additional nuclei beyond two. The appearance of three or more nuclei within a single RPE cell may result from the displacement of nuclei from adjacent cells moving above or below the focal plane. This phenomenon, coupled with parallax effects inherent in standard *en face* imaging techniques, can create the illusion of multinucleation within individual RPE cells. Alternatively, pathological circumstances might lead to cell fusion, which may offer cells the opportunity to share resources and thus save the remaining damaged cells. Cell fusion may also be a potential mechanism that could enable thinner cells to cover more of Bruch’s membrane via an increase in surface area without requiring an increase in cell number. There is good evidence to support this model in aging and potential stress models (Chen et al., 2016), thus it becomes important to ensure that we accurately identify which nuclei belong to which cell and whether or not multinucleate cells are true hallmarks of abnormal RPE sheet or potential imaging artifact.

A potential confound of visual RPE nuclei localization in standard *en face* images is that the choroid has many nucleated cells. Show-through of endothelial cells can occur if the density of pigment granules of the RPE or melanocytes is low, which might occur in disease and therefore it is essential that we do not mistakenly count any of these nuclei when determining RPE nuclei counts per cell. Thus, it is critical to accurately observe the location and shape of nuclei in order to establish whether a given nucleus is really a nucleus of the RPE or another confounding layer in a flatmount.

Standard imaging of *ex vivo* RPE sheet preparations typically involves simultaneous immunofluorescent staining of apical cell border proteins, such as ZO-1, and nuclei with different wavelength fluorophores. Images acquired with a confocal microscope are usually at a lower magnification (10-20x objective) as global patterning is of initial interest in multifield stitched images. In some experiments, multiple images in the z-axis are acquired (z-stacks) which are then converted to 2D maximum intensity projections (MIPs) by collapsing the z-axis and forcing a straight apical to basal orientation of RPE cells. This process can introduce the aforementioned parallax as nuclei from adjacent cells can actually appear within the cell border of a central cell if enlarged at only the apical surface, and not uniformly throughout the z-axis. While some 3D ultrastructure imaging of RPE cells has been achieved via serial block face scanning electron microscopy (SBF-SEM)(Keeling et al., 2020), this method still presents a technical barrier for many labs as well as not being suited for simultaneous whole eye image acquisition used in disease phenotyping.

A high magnification (60x) 3D rendering of the RPE sheet with fluorescent staining of the apical cell borders, basal, and lateral faces of individual RPE cells can help us decode patterns associated with normal RPE cell properties, as well as those in stressed or damaged conditions. Additionally, when imaged with modern, automated confocal microscopes, entire eye flatmount preparations can be obtained for large scale pattern recognition before subsequent imaging of specific dysmorphic cells of interest.

Thoughtful interpretation of 3D ultrastructure imaging should also be applied to other RPE cell dysmorphia. We and others find that RPE cells undergo progressive degeneration in patients (Bhatia et al., 2016b; Rashid et al., 2016; Wang et al., 2018; Q. Zhang et al., 2019) and in animal models of retinal damage or disease (Cano et al., 2014; Chrenek et al., 2012; N. Zhang et al., 2021), exhibiting characteristics of dedifferentiation reminiscent of epithelial-to-mesenchymal transition (EMT; reviewed in (Datta et al., 2017). Experiments with cultures of RPE cells or explants also show that when stressed, RPE cells appear to dedifferentiate via EMT-like processes (Cano et al., 2014; Datta et al., 2017, 2023; Mertz et al., 2021; Sripathi et al., 2023; Sripathi, Hu, Liu, et al., 2021; Sripathi, Hu, Turaga, et al., 2021)(22,54-59).

The existence of RPE “epithelial-to-mesenchymal transition” (RPE EMT) may imply that RPE cells lose apical-basal polarity, normal intercellular adhesion, and outer blood-retina barrier functions. RPE cells might become mesenchymal cells because they have begun to move and migrate and invade the neural retina. This transition might include the loss of E- and P-cadherin and increase expression of putative “mesenchymal markers” including SMA, Vimentin, Snail, Twist, Slug. Here we image complete RPE monolayers after NaIO_3_ tail vein injections, illustrating stress responses of RPE cells in different locations demonstrating variable pleomorphic and polymorphic responses that differ radially. The degree of response is also indicated by apparent changes in activity of a CAG promoter driving ZsGreen expression consistent with increased activity reminiscent of changing from an epithelial level of low expression to a higher level of activity like that found in mesenchymal cells (Caminos et al., 2014).

“RPE EMT” may represent a wound-healing response, potentially involving fibrosis, triggered by external signals such as growth factors, cytokines, and extracellular matrix components. While it remains uncertain whether “RPE EMT” constitutes a true epithelial-mesenchymal transition, it likely shares commonalities with other EMT processes, suggesting that therapies targeting EMT in other contexts could be effective for RPE-related diseases and wound healing.

## Methods

### Animal models of RPE damage

All mouse handling procedures and care were approved by the Emory Institutional Animal Care and Use Committee and followed the ARVO Statement for the Use of Animals in Ophthalmic and Vision Research. Adult (postnatal day 90) male C57BL6/J mice were obtained from The Jackson Laboratory (Bar Harbor, ME) and were housed under a 12:12-hour light–dark cycle. Following our previously published protocol (Chrenek et al., 2016), one mouse was exposed to toxic levels of light which caused light-induced retinal degeneration (LIRD). In a second mouse, a subretinal injection surgery was performed, causing a temporary retinal detachment and subsequent RPE sheet disruption (Donaldson et al., 2018). Both animals were sacrificed and tissue harvested a few days after the respective procedures. Tail-vein injections were conducted as previously described (N. Zhang et al., 2021). We employed an inducible VMD2-Cre driver line(Le et al., 2008) (JAX strain# 032910) crossed to a CAG-Lox-Stop-Lox-ZsGreen mouse line (Madisen et al., 2010) (JAX strain# 007906) all on the C57BL/6J background. This cross yielded RPE cells that were indelibly tagged with ZsGreen expression. Any cells that were originally normal, mature RPE in phenotype and subsequently “transdifferentiated” into another cell type remain permanently marked by ZsGreen fluorescence, regardless of their phenotypic fate

### Tissue preparation and immunofluorescent labeling

Whole eye RPE flatmounts (FMs) were prepared following our previously published protocols (Boatright et al., 2015; Jiang et al., 2013) with some minor modifications (**Figs 1, 4, 5; Vid 1 and 2**). Eyes were fixed in Z-Fix (Anatech Ltd, Battle Creek, MI, USA) for 10 min, and then washed three times with Hank’s Balanced Salt Solution (HBSS; Cat. #14025092, Gibco by Life Technologies, Grand Island, NY, USA). Eyes were stored at 4°C for up to 24 hours before being dissected. Following the removal of the iris and neural retina, four radial cuts were made to produce four RPE–scleral flaps with additional smaller relief cuts to aid with flattening of the tissue. FMs were then mounted RPE side up, on conventional microscope slides to which a silicon gasket had been applied (Grace Bio-Labs, Bend, OR, USA).

**Figure 1.**
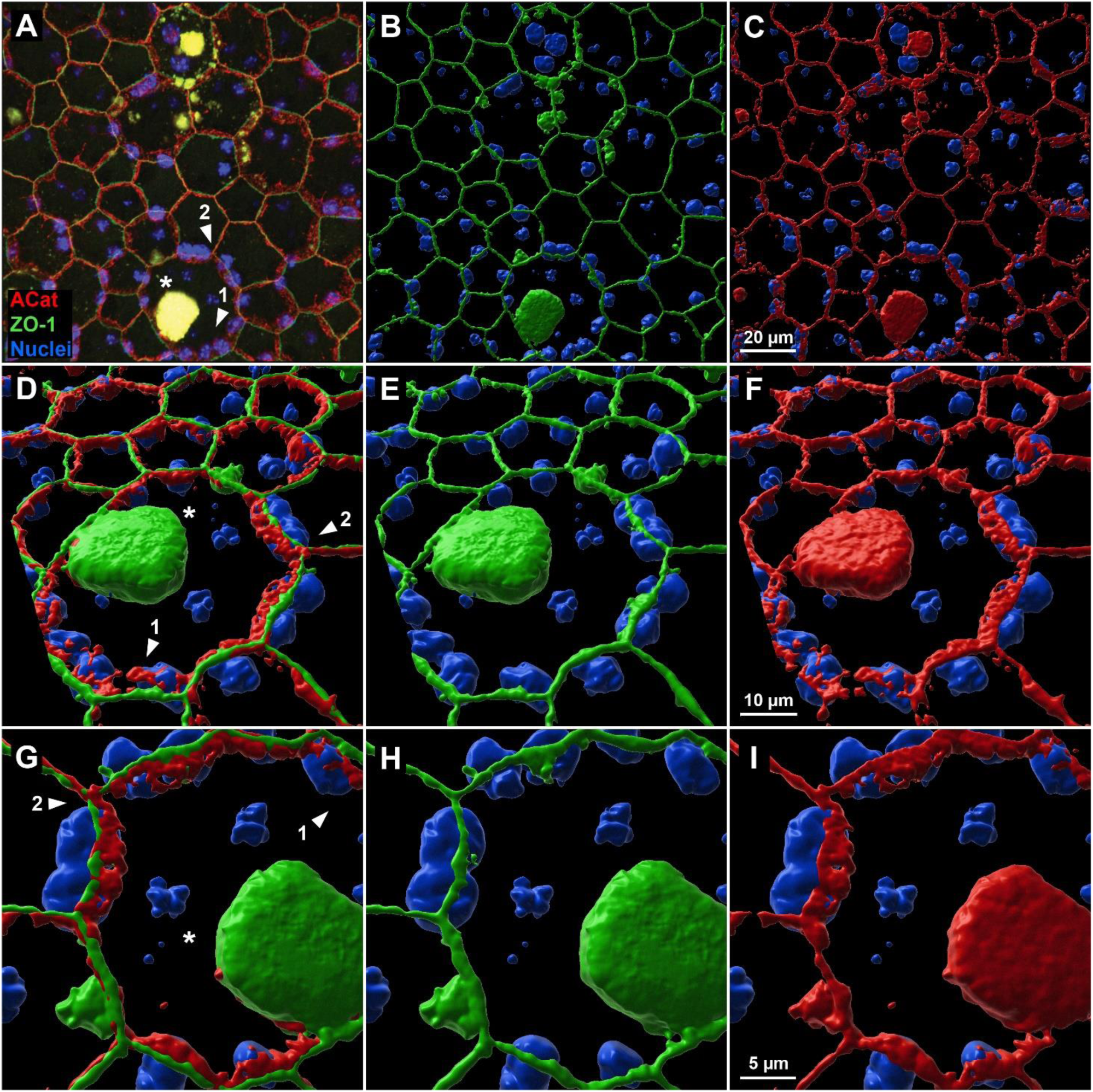
3D reconstruction of apical and lateral structure of single cells in RPE sheet following LIRD. **(A)** Original ROI selection from entire RPE-FM confocal image with labeling of cell perimeter marker ZO-1 (green), lateral surface and perimeter marker alpha-catenin (red) and nuclei (blue). 3D reconstructions of the same ROI were made using IMARIS to show nuclear position in relation to ZO-1 **(B, E, H)** and alpha-catenin **(C, F, I)** separately as well as combined **(D,G)**. Numbered arrows **(1, 2)** reference multinuclear clusters of interest which are then further visualized at increasing magnification. **Arrow 1** highlights an abnormally large RPE cell with apparent multinucleation in original 2D confocal image **(A)**. Higher magnification and alternate viewing angles reveal instead that multiple nuclei are in highly lateralized positions of adjacent cells (also seen near **Arrow 2**). **Asterisk(*)** indicates auto fluorescent aggregate frequently found in LIRD-affected RPE tissue. **Video 1** provides additional viewing angles and magnification of this specific ROI.

The FMs were rinsed with HBSS, then incubated in blocking buffer consisting of 1% BSA (Sigma, St. Louis, MO, USA) in 0.1% Triton X-100 (Sigma) HBSS solution for 1 hour at room temperature. Primary incubation occurred overnight at room temperature (1:100 anti-ZO-1, Cat. #MABT11, MilliporeSigma; 1:500 anti-CTNNA1 (alpha-catenin)[EP1793Y], Cat. #ab51032, Abcam, Cambridge, MA, USA). Following washes with 0.1% Triton-X-100 in HBSS, FMs were incubated with secondary antibodies (Alexa Fluor 488,1:1000 donkey anti-rat immunoglobulin G, Cat. #A21208, Thermo Fisher Scientific, Waltham, MA, USA; Alexa Fluor 568, 1:1000 goat anti-rabbit immunoglobulin G, Cat. #A11036, Thermo Fisher Scientific) overnight at room temperature. FMs were washed with Hoechst 33258 nuclear stain in 0.1% Triton X-100 in HBSS three times, followed by two washes with only wash buffer, and then mounted with Fluoromount-G (Cat. #17984-25, Electron Microscopy Sciences, Signal Hill, CA, USA), coverslipped, and allowed to set overnight.

Ocular cross sections (**Fig 2, Vid 3**) were prepared using a modification of the freeze substitution technique described by Sun *et al* (Sun et al., 2015). Specifically, after enucleation, eyes were rapidly frozen in 10 ml of 3% acetic acid in methanol that had been prechilled in dry ice. The eyes were fixed and dehydrated at −80°C for 4 days, cleared with methanol and xylenes, and embedded in paraffin. 5 um sections were cut and immunostained with anti-alpha-catenin antibody (same antibody as FM preparation), counterstained with Hoechst 33342, and imaged with a Nikon Ti2 microscope with A1R confocal imager with a 40x silicone oil objective with 2.5x zoom.

**Figure 2.**
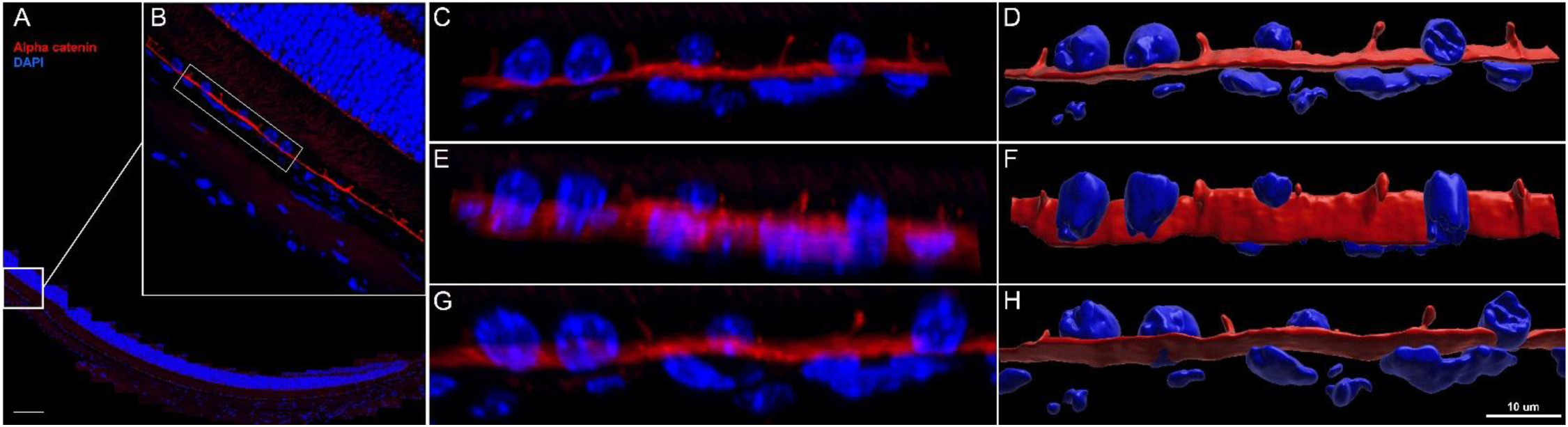
Confocal images and corresponding 3D reconstruction of the retina-RPE-choroid with lateral and basal surface labeling of RPE cell membranes in normal mouse eyes. **(A,B)** Region of interest (ROI) selection from a 5 µm-thick cross-section labeled for alpha-catenin (red) and nuclei (blue), scale bar = 100 µm. A 3-cell-wide region (bounding box in **B**) was isolated to provide alternate-angle views **(C, E, G)** and corresponding 3D reconstructions **(D, F, H)**. C and D are lateral views showing distinct morphologies in RPE nuclei (5 µm, round/potato shapes, 5-6 speckles) versus choriocapillaris nuclei (axially thin, irregular, less speckling). Alpha-catenin labels lateral and basal faces with uniform staining and thickness. Lateral RPE walls show no clear orientation pattern in adjacent cells with angling > 45° relative to basal surfaces (closer to perpendicular than not). **Video 3** of reconstructed ROI rotated for viewing from multiple angles corresponds to **D**, **F**, and **H**.

### Confocal microscopy and 3D image visualization

Whole-eye FMs were imaged in their entirety at lower magnification (10x objective lens) using a Nikon Ti microscope with C1 or A1R confocal scanner (Nikon Instruments Inc., Melville, NY, USA), with sufficient z-stack thickness to capture entire RPE cells throughout the FM. Regions of interest (ROIs) containing dysmorphic cells were then reimaged at higher magnification (60x and 100x objective lenses) with appropriate z-slice spacing to satisfy the Nyquist requirement for 3D capture and viewing. Imaris (version 10.1, Bitplane, Oxford, UK) software was used to isolate and post-process ROIs from raw confocal files as well as generate videos highlighting cell membrane, internal alpha-catenin structure, and nuclear position from multiple viewing angles.

## Results

We present images, reconstructions, and videos generated from high magnification confocal z-stacks of two damage-model RPE sheets highlighting 3D ultrastructure of individual cells. Specifically, we selected ROIs that exhibit a heterogenous mix of normal and dysmorphic cells and include larger than normal, multinucleate cells (**Fig 1**, **Vid 1** and **2**). Extending the visualization of conventionally sectioned ocular tissue, we also present 3D images, reconstructions and videos of lateral and basal RPE cell surfaces (**Fig 2**, **Vid 3**). Finally, we show both local and global disruptions in the RPE sheet reminiscent of EMT in entire mouse eyes following induced RPE loss via intravenous tail vein NaIO_3_ injections (**Fig 4** and **5**).

**Figure 4.**
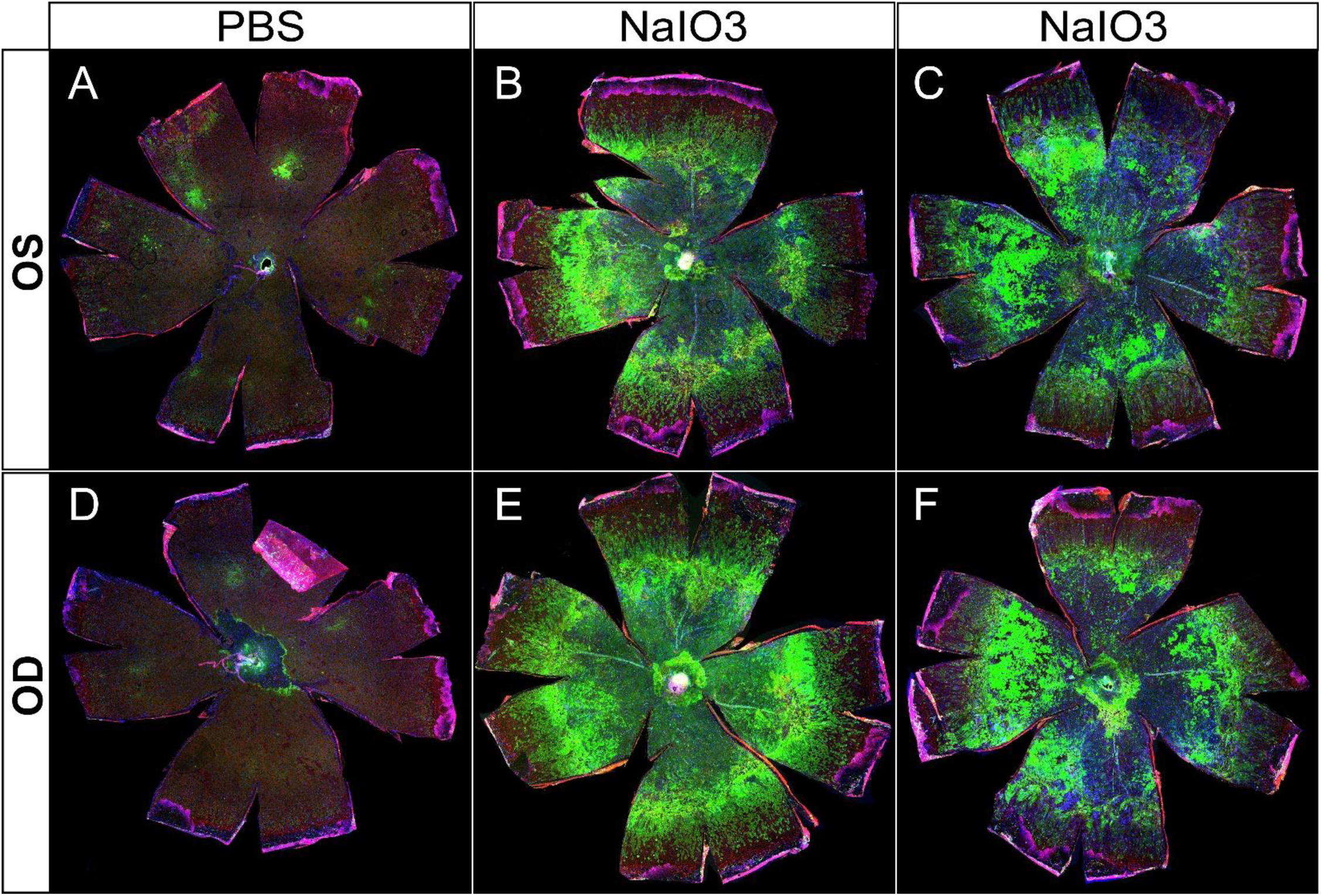
Representative examples of whole-eye flatmounts prepared 8 days following control (PBS) or NaIO_3_ tail vein injections. Drastically altered RPE morphology was observed in both eyes of animals following intravenous injection of NaIO_3_ (**B** and **E**, **C** and **F**) but not in those injected with only PBS vehicle (**A** and **D**). Of note is a general gradient of cellular stress severity, originating from the optic nerve head. A large central atrophic lesion is surrounded by a ring-like border region composed of cellular debris and aggregates (bright green = ZsGreen expression driven by CAG promoter solely in RPE cells or descendants), most likely dying RPE cells. Continuing radially, an EMT-like response is seen characterized by elongation of RPE cells before transitioning to unaffected peripheral cells.

**Figure 5.**
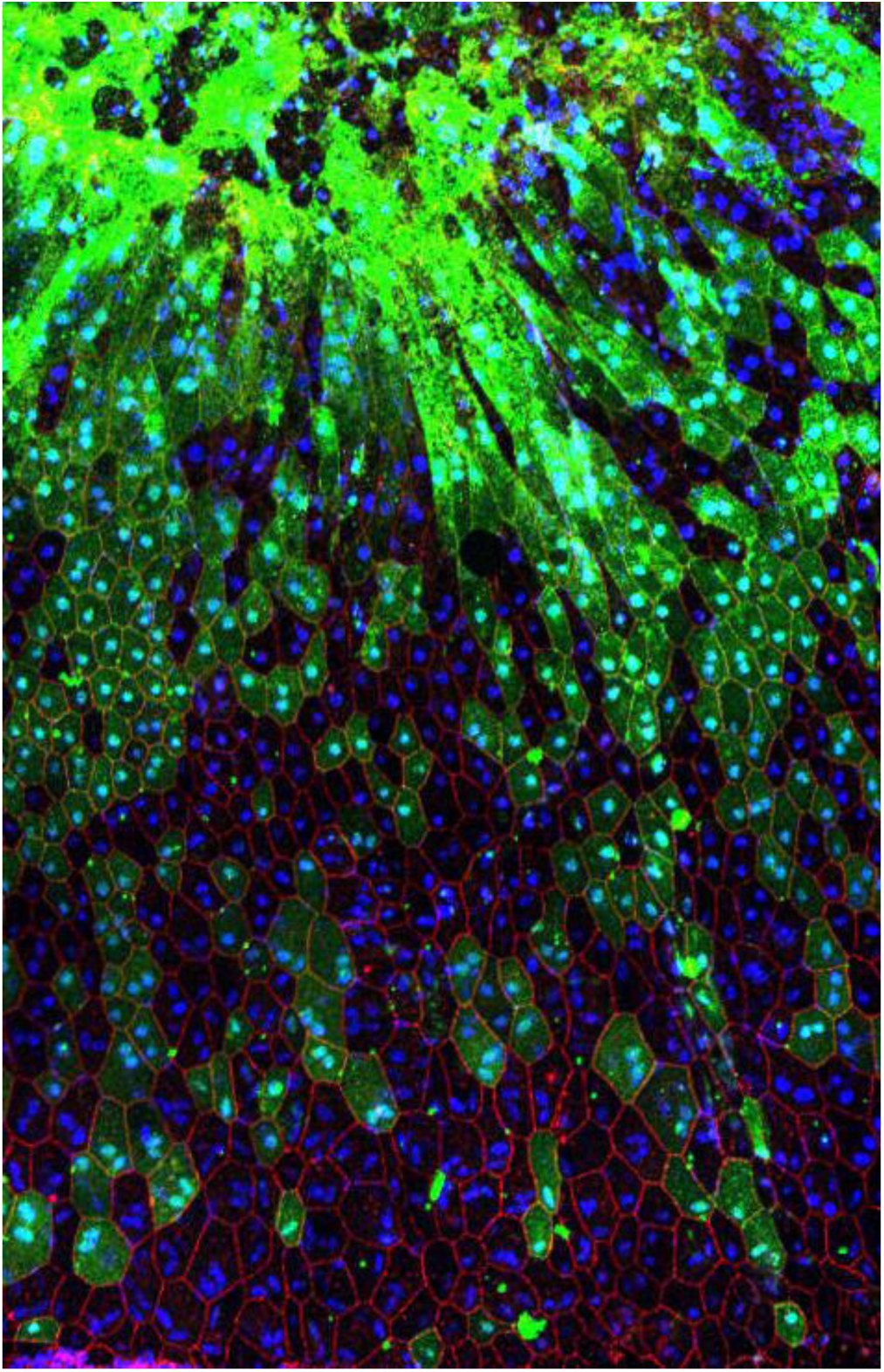
Region of Interest of “RPE-EMT” transitional region following NaIO_3_ administration. Extremely high expression of ZsGreen linked to CAG promoter activity is evident in regions where cell morphology is severely abnormal (bright green top 1/3 of image) and apical cell border delineation is non-existent (red, ZO-1). Adjacent, intact cells are elongated before transitioning to more regular morphology further from the atrophic area.

**Video 1** presents an isolated z-stack imaged at 100x magnification with labeling of ZO-1 (green), alpha-catenin (red), and nuclei (blue) from a LIRD-damaged whole-eye RPE FM. Initially oriented basal-side up, with the apical surface away from the viewer, a large, seemingly multinucleate cell is found in the top-middle of the image. The cell is much larger than normal by about 3-fold on each of the X- and Y-axes, suggesting an 8-10-fold increase in cell size. This cell has a number of nuclei around its perimeter that appear to be within the cell depending on viewing angle. Additionally, there is a large auto-fluorescent aggregate (appearing as yellow here) of unknown composition, which is hypothesized to be internalized rod/cone outer segments or lipofuscin, contained within the cell. Other cells of interest include those at the bottom middle, where again it appears that larger cells are multinucleated with a number of nuclei appearing near their perimeters. As playback proceeds, orientation is flipped with the apical surface being at the top when viewed orthogonally and closest to the viewer later when viewed *en face*. The viewer is then “flown” through the image where the auto fluorescent aggregate and laterally oriented alpha-catenin is visible below the apically located ZO-1 rim. Zooming closer and viewing from the apical surface, reveals multiple nuclei clustered around the perimeter of the enlarged cell. The alpha-catenin signal is removed to highlight the ZO-1 outline, mimicking typical RPE cell border delineation imaging. When the alpha-catenin channel is restored, it is apparent that the large cell is overhanging these nuclei on the ring and edge. When rotated to a side or underneath view, it becomes clear that these nuclei actually belong not to the central cell, but to the adjacent neighboring cells, and the adjacent cells have been pushed outward. We do not know yet whether it is the central large cell pushing over the top of the adjacent neighbors or if the neighbors are pulling on the central cell resulting in over-coverage of the neighboring cells.

**Video 2** presents a 3D view of a region which contains greatly dysmorphic RPE cells following SRI-induced retinal detachment and recovery. Here, relying on ZO-1 (green) solely for individual cell demarcation (as is typically found in the field) is problematic as abnormally large cells are adjacent to select cells of strikingly smaller area. This “purse-string” pattern has been observed in multiple models of RPE stress when viewed in *en face* 2D images and potentially reflects cells in the process of programmed cell death and/or removal from the sheet. However, it is unclear if the larger cells are actively pushing out the smaller cells or merely expanding to fill in the previously occupied space to maintain the RPE’s barrier integrity. When imaged with concurrent alpha-catenin staining (red), the viewer can observe many different underlying cellular shapes that don’t directly align with the ZO-1 perimeter staining. For instance, in the top middle of the video, there is a large, spherical structure of labeled alpha-catenin that spreads underneath multiple ZO-1 delineated perimeters. This structure encompasses two nuclei (blue), separating them from the pair of nuclei directly below. Without this additional ultrastructural information, the viewer might erroneously conclude that all four nuclei are contained in an enlarged, multinucleate cell, when in reality it is two adjacent cells. When looking at the enlarged cell in the right-middle region of the video (just below the ZO-1 labeled cell of very small area), the viewer appears to see four nuclei encompassed by the ZO-1 perimeter. However, when viewed as a 3D aspect, it is apparent that the left pair of nuclei are visible through an open cylinder of alpha-catenin, suggesting that they are located in a different cell, with the pair on the right more superficially separated. Special care should be taken to determine in 3D space if these two pair of nuclei are contained by the enlarged cell or are indeed in separate cells. Thus, when viewed in 3D with the addition of lateral labeling, this video exemplifies the multitude of dynamic and dysmorphic reorganization that occurs following RPE stress.

In **Figures 4 and 5**, we present images from untreated and NaIO_3_ stressed mouse RPE flat mounts. Green fluorescence indicates RPE cells expressing ZsGreen in generally high levels, partly masked in untreated RPE by the heavy melanin of the C57Bl/6J strain. The mice used in these experiments were genetically tagged using a floxed mouse line crossed to an RPE specific Cre driver line. The resulting cross yielded mice with a permanent genetic marker of RPE cells. Any RPE cell that expressed Cre from the Best1 Cre driver line was tagged as green because of the Cre driven deletion of the Stop codon from the CAG-lox-stop-lox-zsGreen flox construct. While not all RPE cells expressed sufficient Cre, about 75-90 percent of the RPE cells did, rendering those cells as green, and that tag was permanent because of the floxed gene in the endogenous genome of the given cell. Even if that RPE cell divided, the resulting daughter cells would still be marked as green, and likewise, if the green RPE cell transitioned to any other phenotype or cell type, that cell would still remain tagged by heavy expression from the permanently encoded endogenous CAG-ZsGreen transgene in that cell’s genome. While the CAG promoter is active in virtually all cell types, it is especially active in motile, mesenchymal cell types. Other retinal cell types express different CAG promoter activity in different cell types (Caminos, Vaquero, Garcia-Olmo, 2014), but the very bright ZsGreen protein still makes almost all cell types patently green by fluorescence microscopy of any sort.

In Figure 4, we illustrate a series of RPE flatmounts from untreated (**4A, 4D**) and NaIO_3_ treated mice (**4B, 4C, 4E, and 4F**). The ZsGreen in **4A** and **4B** was partly masked by heavy melanin pigment normally found in C57BL/6J mice, but the RPE cells do display ZsGreen fluorescence. After NaIO_3_ oxidative stress, many RPE cells were killed in the central posterior third to half of the flatmounts (Fig. panels **4B, 4C, 4E, and 4F**). Only some residual cell debris retained any ZsGreen fluorescence. In a mid-periphery bullseye ring, there was heavy ZsGreen fluorescence coincident with highly elongated and irregularly shaped cells that appeared to pile on top of each other. Farther more peripherally there was a bullseye zone of elongated cells with distinct ZO-1 outlines suggesting that these cells retained the characteristic RPE apical basal polarity, cell-to-cell contact, and adherence in a single monolayer to underlying Bruch’s membrane even though elongated radially. Even farther out into the periphery to the ciliary body, the RPE cells appeared to exhibit normal shape and size, characteristic of that far peripheral zone (Kim et al., 2021).

In **Figure 5**, we illustrate a closeup of the cells from a NaIO_3_ treated mouse. The highly elongated phenotype is evident in the close-up. Cells that expressed more green fluorescence appeared to be more elongated. The more elongated the cell, the greater to loss of ZO-1 staining and the greater the loss of straight edges, symmetry, and regularity. It is more difficult to determine the location of nuclei in cells as the cells became highly elongated.

## Conclusions

The images and work presented here emphasize the need to determine the location of nuclei accurately, precisely, and reliably in RPE cells. By using relatively standard confocal microscopy but with the appropriate approach to instrumentation and image acquisition settings, we simply employ a careful examination of nuclear localization in 3D. We used Imaris, a user-friendly, commercially available program for visualization and creation of videos, however other software (including ImageJ with appropriate plugins) would likely be adequate for the same purpose.

A key question when analyzing RPE dysmorphia is if neighboring cell nuclei remain in their respective origin cells but are visualized within an enlarged central cell’s borders when viewed *en face*, or are indeed components of a larger multinucleate cell. Our results suggest that when looking at large, multinuclear cells in a RPE sheet which has been subjected to stress, it is important to visualize these changes from a 3D perspective at the individual cell level. In multiple images, we show that nuclei which appear as members of a large multinucleate cell are actually still members of surrounding cells when viewed at higher magnification and alternate angles than the standard apical to basal collapsed view.

Correct interpretation of multiple nuclei in RPE cells depends on a better understanding of how the basal and lateral faces of the RPE cells move following aging and stress. A small proportion of RPE cells appear capable of greatly increasing in size following severe stress. Largely absent under normal conditions, it remains unclear if all RPE cells have this capability to vastly enlarge, or if only a small subclass of RPE cells can do this. Further experiments investigating damage-associated morphological abnormalities are required to understand the unusually large cell phenotype and find potential common traits, if any, between damage conditions. It is likewise unclear if these cells can undergo further morphological or functional transformations such as, but not limited to, epithelial–mesenchymal transition (EMT).

Finally, a fate-mapping strategy employing RPE cells genetically marked by a Cre-lox system allows accurate tracing of the lineages of cells of previously unknown origins in the subretinal space. We were able to indelibly mark cells of extremely elongated and irregular shapes and divine that they were of RPE origins with this strategy in mice treated by tail vein injection of small amounts of NaIO_3_.

## Supporting information

Vid 1: 3D LIRD-induced RPE dysmorphia

Vid 2: 3D SRI-induced RPE dysmorphia

Vid 3: 3D RPE histological section

Fig 4A: Full RPEFM-PBS-OS

Fig 4D: Full RPEFM-PBS-OD

Fig 4B: Full RPEFM-NaIO3-OS

Fig 4E: Full RPEFM-NaIO3-OD

Fig 4C: Full RPEFM-NaIO3-OS

Fig 4F: Full RPEFM-NaIO3-OD

---

**Video 1.**
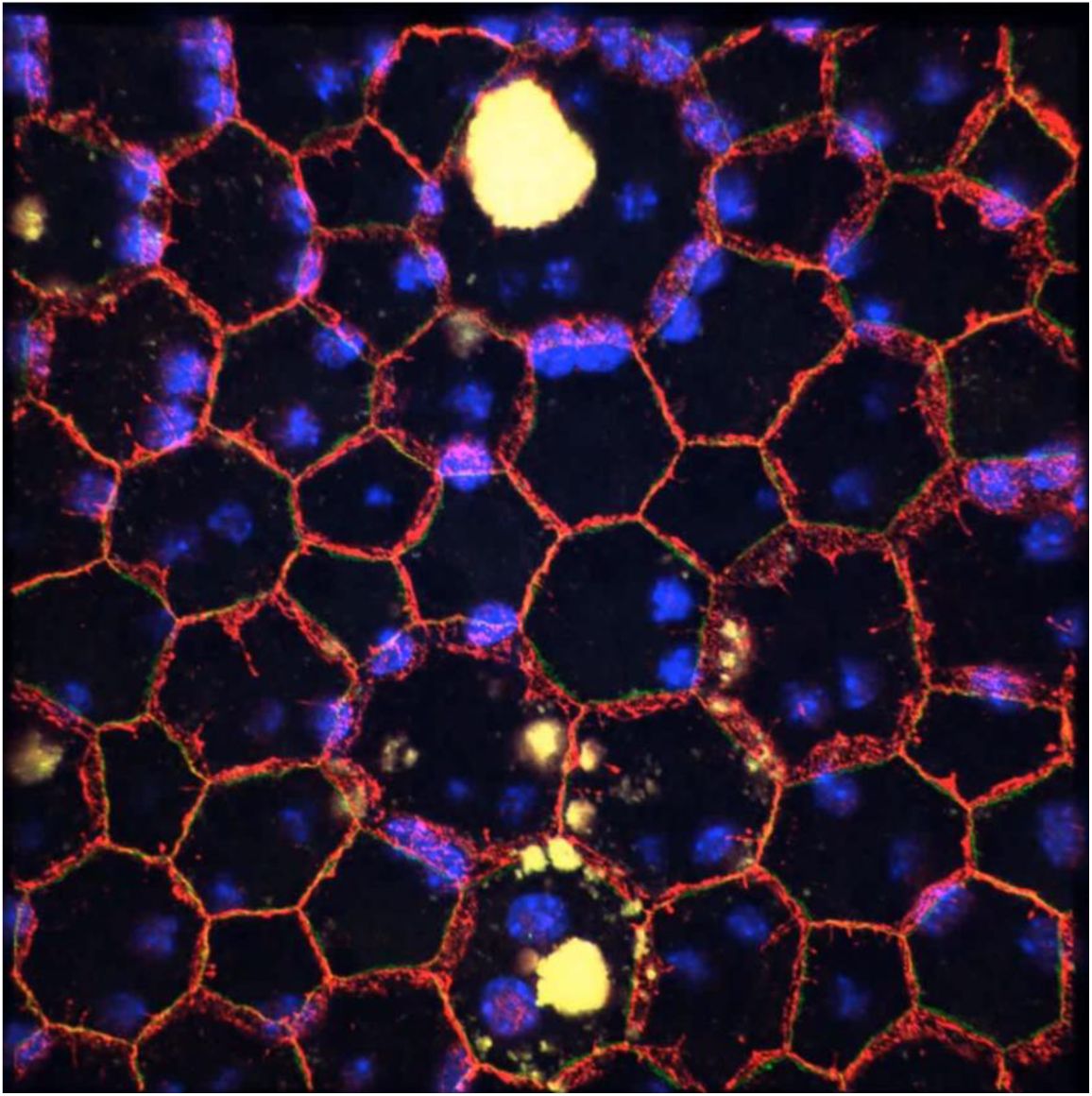
Multi-angle and magnification viewing of LIRD-affected RPE cells. 3D video of confocal z-stack (100x magnification) showing ZO-1 cell perimeter (green), alpha-catenin lateral and perimeter (red) and nuclear (blue) labeling from various angles not available in traditional 2D *en face* viewing from the apical surface. RPE cells have developed dysmorphia following LIRD. Large, seemingly multinucleate cells are shown to contain only a maximum of two nuclei when viewed from alternate angles and with concurrent lateral labeling of alpha-catenin.

**Video 2.**
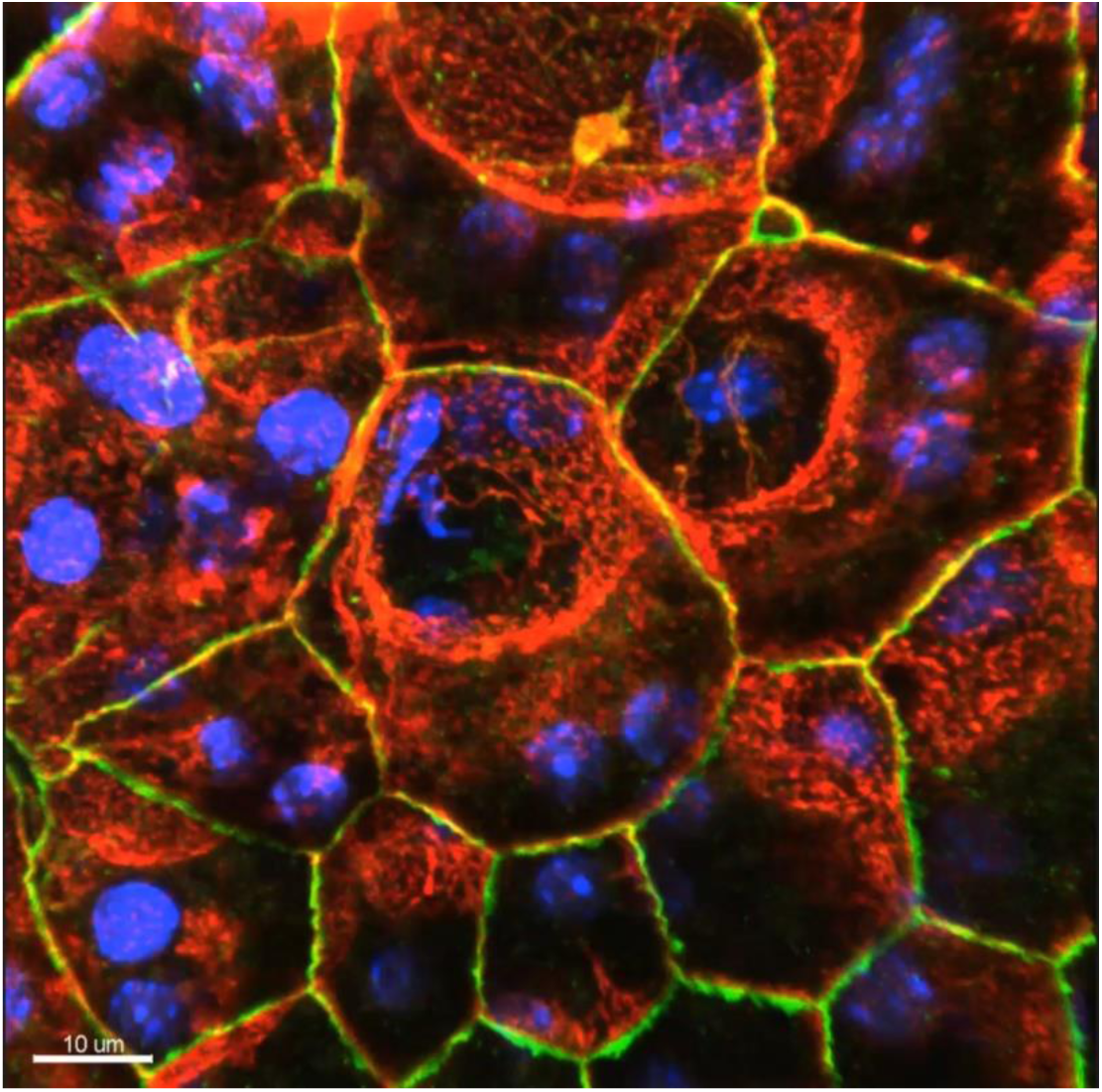
Multi-angle view of RPE cell dysmorphia following SRI-induced retinal detachment. 3D video of confocal z-stack (100x magnification) showing ZO-1 cell perimeter (green), alpha-catenin lateral and perimeter (red) and nuclear (blue) labeling. Vast changes in individual RPE cell size, shape, orientation and alignment are visible during recovery from SRI-induced retinal detachment. Again, large, seemingly multinucleate cells are shown to contain less nuclei than expected when counted using ZO-1 perimeter staining and typical apical viewing, instead of lateral labeling of cell membranes and alternative viewing angles in 3D.

**Video 3.**
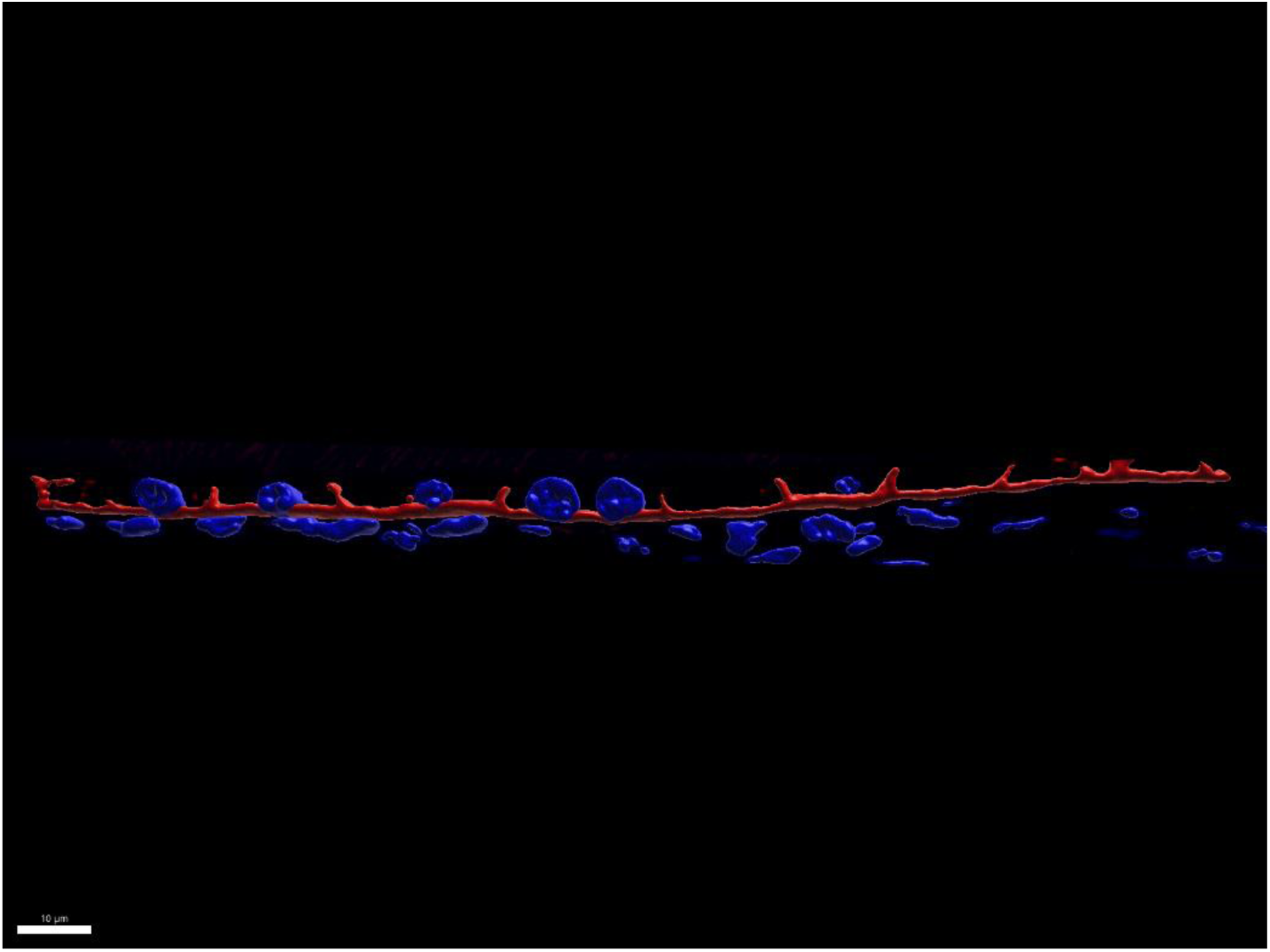
Rotation of 3D reconstruction selected from conventional cross sections for alternate angle viewing of lateral and basal face labeling. Initially starts with single FOV highlighted in **Fig 2B** before zooming into selected region of interest (ROI) from **Fig 2C-H** and then rotated through 3D space. ROI is then widened to view entire lateral portion of RPE/choroid interface, highlighting differences in nuclear structure and position between RPE and underlying choriocapillaris.

## References

Bhatia, S. K., Rashid, A., Chrenek, M. A., Zhang, Q., Bruce, B. B., Klein, M., Boatright, J. H., Jiang, Y., Grossniklaus, H. E., & Nickerson, J. M. (2016a). Analysis of RPE morphometry in human eyes. Molecular Vision, 22, 898–916.

Bhatia, S. K., Rashid, A., Chrenek, M. A., Zhang, Q., Bruce, B. B., Klein, M., Boatright, J. H., Jiang, Y., Grossniklaus, H. E., & Nickerson, J. M. (2016b). Analysis of RPE morphometry in human eyes. Molecular Vision, 22, 898–916.

Boatright, J. H., Dalal, N., Chrenek, M. A., Gardner, C., Ziesel, A., Jiang, Y., Grossniklaus, H. E., & Nickerson, J. M. (2015). Methodologies for analysis of patterning in the mouse RPE sheet. Molecular Vision, 21, 40–60.

Bonilha, V. L. (2008). Age and disease-related structural changes in the retinal pigment epithelium. Clinical Ophthalmology, 2(2), 413–424. 10.2147/opth.s2151

Caminos, E., Vaquero, C. F., & García-Olmo, D. C. (2014). “Green mice” display limitations in enhanced green fluorescent protein expression in retina and optic nerve cells. Histology and Histopathology, 29(12), 1601–1612. 10.14670/HH-29.1601

Cano, M., Wang, L., Wan, J., Barnett, B. P., Ebrahimi, K., Qian, J., & Handa, J. T. (2014). Oxidative stress induces mitochondrial dysfunction and a protective unfolded protein response in RPE cells. Free Radical Biology & Medicine, 69, 1–14. 10.1016/j.freeradbiomed.2014.01.004

Chen, M., Rajapakse, D., Fraczek, M., Luo, C., Forrester, J. V., & Xu, H. (2016). Retinal pigment epithelial cell multinucleation in the aging eye – a mechanism to repair damage and maintain homoeostasis. Aging Cell, 15(3), 436. 10.1111/acel.12447

Chrenek, M. A., Dalal, N., Gardner, C., Grossniklaus, H., Jiang, Y., Boatright, J. H., & Nickerson, J. M. (2012). Analysis of the RPE sheet in the rd10 retinal degeneration model. Advances in Experimental Medicine and Biology, 723, 641–647. 10.1007/978-1-4614-0631-0_81

Chrenek, M. A., Sellers, J. T., Lawson, E. C., Cunha, P. P., Johnson, J. L., Girardot, P. E., Kendall, C., Han, M. K., Hanif, A., Ciavatta, V. T., Gogniat, M. A., Nickerson, J. M., Pardue, M. T., & Boatright, J. H. (2016). Exercise and Cyclic Light Preconditioning Protect Against Light-Induced Retinal Degeneration and Evoke Similar Gene Expression Patterns. Advances in Experimental Medicine and Biology, 854, 443–448. 10.1007/978-3-319-17121-0_59

Christensen, D. R. G., Brown, F. E., Cree, A. J., Ratnayaka, J. A., & Lotery, A. J. (2017). Sorsby fundus dystrophy – A review of pathology and disease mechanisms. Experimental Eye Research, 165, 35–46. 10.1016/j.exer.2017.08.014

Datta, S., Cano, M., Ebrahimi, K., Wang, L., & Handa, J. T. (2017). The impact of oxidative stress and inflammation on RPE degeneration in non-neovascular AMD. Progress in Retinal and Eye Research, 60, 201–218. 10.1016/j.preteyeres.2017.03.002

Datta, S., Cano, M., Satyanarayana, G., Liu, T., Wang, L., Wang, J., Cheng, J., Itoh, K., Sharma, A., Bhutto, I., Kannan, R., Qian, J., Sinha, D., & Handa, J. T. (2023). Mitophagy initiates retrograde mitochondrial-nuclear signaling to guide retinal pigment cell heterogeneity. Autophagy, 19(3), 966–983. 10.1080/15548627.2022.2109286

Donaldson, K. J., Skelton, H. M., Sellers, J. T., Grossniklaus, H., & Nickerson, J. M. (2018). Novel ex and in vivo methods for non-invasive longitudinal tracking of RPE dysmorphology following subretinal injections. Investigative Ophthalmology & Visual Science, 59(9), 4982.

Gambril, J. A., Sloan, K. R., Swain, T. A., Huisingh, C., Zarubina, A. V., Messinger, J. D., Ach, T., & Curcio, C. A. (2019). Quantifying Retinal Pigment Epithelium Dysmorphia and Loss of Histologic Autofluorescence in Age-Related Macular Degeneration. Investigative Ophthalmology & Visual Science, 60(7), 2481–2493. 10.1167/iovs.19-26949

Hergott, G. J., Sandig, M., & Kalnins, V. I. (1989). Cytoskeletal organization of migrating retinal pigment epithelial cells during wound healing in organ culture. Cell Motility and the Cytoskeleton, 13(2), 83–93. 10.1002/cm.970130203

Jiang, Y., Qi, X., Chrenek, M. A., Gardner, C., Boatright, J. H., Grossniklaus, H. E., & Nickerson, J. M. (2013). Functional Principal Component Analysis Reveals Discriminating Categories of Retinal Pigment Epithelial Morphology in Mice. Investigative Ophthalmology & Visual Science, 54(12), 7274–7283. 10.1167/iovs.13-12450

Kalnins, V. I., Sandig, M., Hergott, G. J., & Nagai, H. (1995). Microfilament organization and wound repair in retinal pigment epithelium. Biochemistry and Cell Biology = Biochimie Et Biologie Cellulaire, 73(9–10), 709–722. 10.1139/o95-079

Keeling, E., Chatelet, D. S., Tan, N. Y. T., Khan, F., Richards, R., Thisainathan, T., Goggin, P., Page, A., Tumbarello, D. A., Lotery, A. J., & Ratnayaka, J. A. (2020). 3D-Reconstructed Retinal Pigment Epithelial Cells Provide Insights into the Anatomy of the Outer Retina. International Journal of Molecular Sciences, 21(21), Article 21. 10.3390/ijms21218408

Kim, Y.-K., Yu, H., Summers, V. R., Donaldson, K. J., Ferdous, S., Shelton, D., Zhang, N., Chrenek, M. A., Jiang, Y., Grossniklaus, H. E., Boatright, J. H., Kong, J., & Nickerson, J. M. (2021). Morphometric Analysis of Retinal Pigment Epithelial Cells From C57BL/6J Mice During Aging. Investigative Ophthalmology & Visual Science, 62(2), 32. 10.1167/iovs.62.2.32

Le, Y.-Z., Zheng, W., Rao, P.-C., Zheng, L., Anderson, R. E., Esumi, N., Zack, D. J., & Zhu, M. (2008). Inducible expression of cre recombinase in the retinal pigmented epithelium. Investigative Ophthalmology & Visual Science, 49(3), 1248–1253. 10.1167/iovs.07-1105

Madisen, L., Zwingman, T. A., Sunkin, S. M., Oh, S. W., Zariwala, H. A., Gu, H., Ng, L. L., Palmiter, R. D., Hawrylycz, M. J., Jones, A. R., Lein, E. S., & Zeng, H. (2010). A robust and high-throughput Cre reporting and characterization system for the whole mouse brain. Nature Neuroscience, 13(1), 133140. 10.1038/nn.2467

Marmor, M. F., & Wolfensberger, T. J. (1998). The retinal pigment epithelium. In Function and Disease (pp. 103–134). https://medtextfree.wordpress.com/2010/12/29/chapter-100-retinal-pigment-epithelium/

Mertz, J. L., Sripathi, S. R., Yang, X., Chen, L., Esumi, N., Zhang, H., & Zack, D. J. (2021). Proteomic and phosphoproteomic analyses identify liver-related signaling in retinal pigment epithelial cells during EMT. Cell Reports, 37(3), 109866. 10.1016/j.celrep.2021.109866

Michaelides, M., Hunt, D. M., & Moore, A. T. (2003). The genetics of inherited macular dystrophies. Journal of Medical Genetics, 40(9), 641–650. 10.1136/jmg.40.9.641

Rashid, A., Bhatia, S. K., Mazzitello, K. I., Chrenek, M. A., Zhang, Q., Boatright, J. H., Grossniklaus, H. E., Jiang, Y., & Nickerson, J. M. (2016). RPE Cell and Sheet Properties in Normal and Diseased Eyes. In C. Bowes Rickman, M. M. LaVail, R. E. Anderson, C. Grimm, J. Hollyfield, & J. Ash (Eds.), Retinal Degenerative Diseases (pp. 757–763). Springer International Publishing. 10.1007/978-3-319-17121-0_101

Sripathi, S. R., Hu, M.-W., Liu, M. M., Wan, J., Cheng, J., Duan, Y., Mertz, J. L., Wahlin, K. J., Maruotti, J., Berlinicke, C. A., Qian, J., & Zack, D. J. (2021). Transcriptome Landscape of Epithelial to Mesenchymal Transition of Human Stem Cell-Derived RPE. Investigative Ophthalmology & Visual Science, 62(4), 1. 10.1167/iovs.62.4.1

Sripathi, S. R., Hu, M.-W., Turaga, R. C., Mertz, J., Liu, M. M., Wan, J., Maruotti, J., Wahlin, K. J., Berlinicke, C. A., Qian, J., & Zack, D. J. (2021). Proteome Landscape of Epithelial-to-Mesenchymal Transition (EMT) of Retinal Pigment Epithelium Shares Commonalities With Malignancy-Associated EMT. Molecular & Cellular Proteomics: MCP, 20, 100131. 10.1016/j.mcpro.2021.100131

Sripathi, S. R., Hu, M.-W., Turaga, R. C., Mikeasky, R., Satyanarayana, G., Cheng, J., Duan, Y., Maruotti, J., Wahlin, K. J., Berlinicke, C. A., Qian, J., Esumi, N., & Zack, D. J. (2023). IKKβ Inhibition Attenuates Epithelial Mesenchymal Transition of Human Stem Cell-Derived Retinal Pigment Epithelium. Cells, 12(8), 1155. 10.3390/cells12081155

Strauss, O. (2005). The retinal pigment epithelium in visual function. Physiological Reviews, 85(3), 845–881. 10.1152/physrev.00021.2004

Sun, N., Shibata, B., Hess, J. F., & FitzGerald, P. G. (2015). An alternative means of retaining ocular structure and improving immunoreactivity for light microscopy studies. Molecular Vision, 21, 428–442.

Wang, J., Zibetti, C., Shang, P., Sripathi, S. R., Zhang, P., Cano, M., Hoang, T., Xia, S., Ji, H., Merbs, S. L., Zack, D. J., Handa, J. T., Sinha, D., Blackshaw, S., & Qian, J. (2018). ATAC-Seq analysis reveals a widespread decrease of chromatin accessibility in age-related macular degeneration. Nature Communications, 9(1), 1364. 10.1038/s41467-018-03856-y

Zhang, N., Zhang, X., Girardot, P. E., Chrenek, M. A., Sellers, J. T., Li, Y., Kim, Y.-K., Summers, V. R., Ferdous, S., Shelton, D. A., Boatright, J. H., & Nickerson, J. M. (2021). Electrophysiologic and Morphologic Strain Differences in a Low-Dose NaIO3-Induced Retinal Pigment Epithelium Damage Model. Translational Vision Science & Technology, 10(8), 10. 10.1167/tvst.10.8.10

Zhang, Q., Chrenek, M. A., Bhatia, S., Rashid, A., Ferdous, S., Donaldson, K. J., Skelton, H., Wu, W., See, T. R. O., Jiang, Y., Dalal, N., & Nickerson, J. M. (2019). Comparison of histologic findings in age-related macular degeneration with RPE flatmount images. Molecular Vision.

