## Supplementary figures and images for "High resolution imaging and interpretation of three-dimensional RPE sheet structure"

### Fig 4A: Full RPEFM-PBS-OS

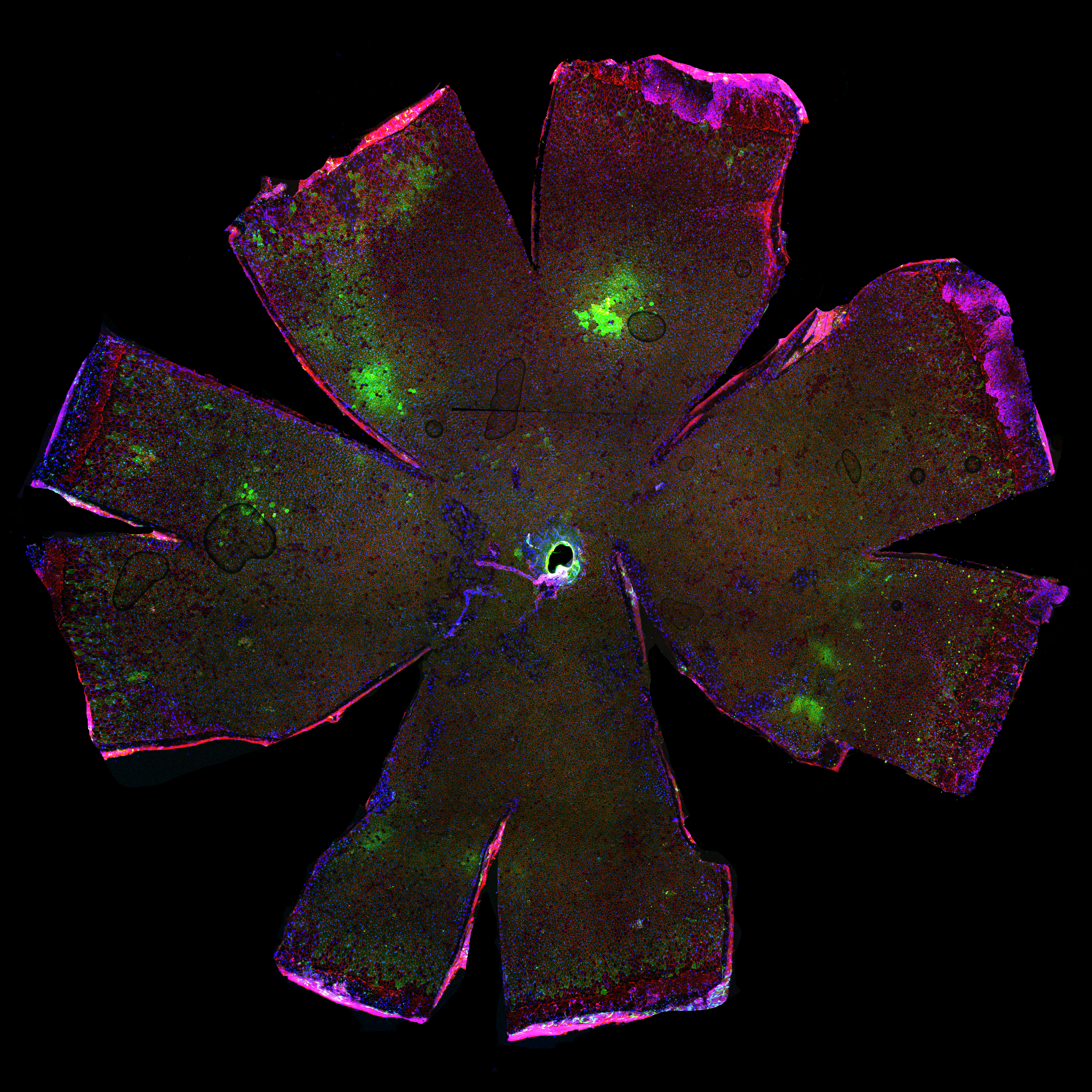

### Fig 4B: Full RPEFM-NaIO3-OS

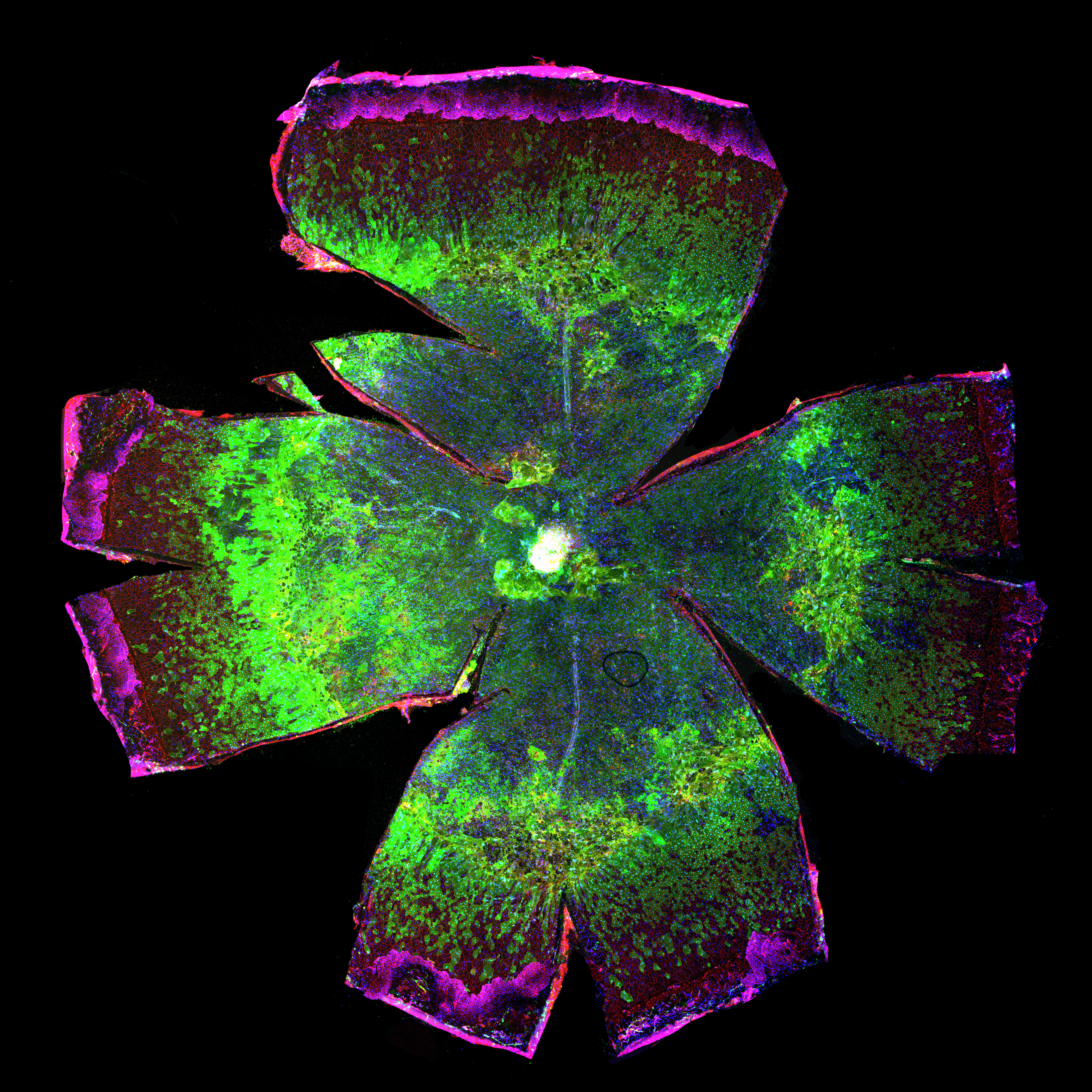

### Fig 4C: Full RPEFM-NaIO3-OS

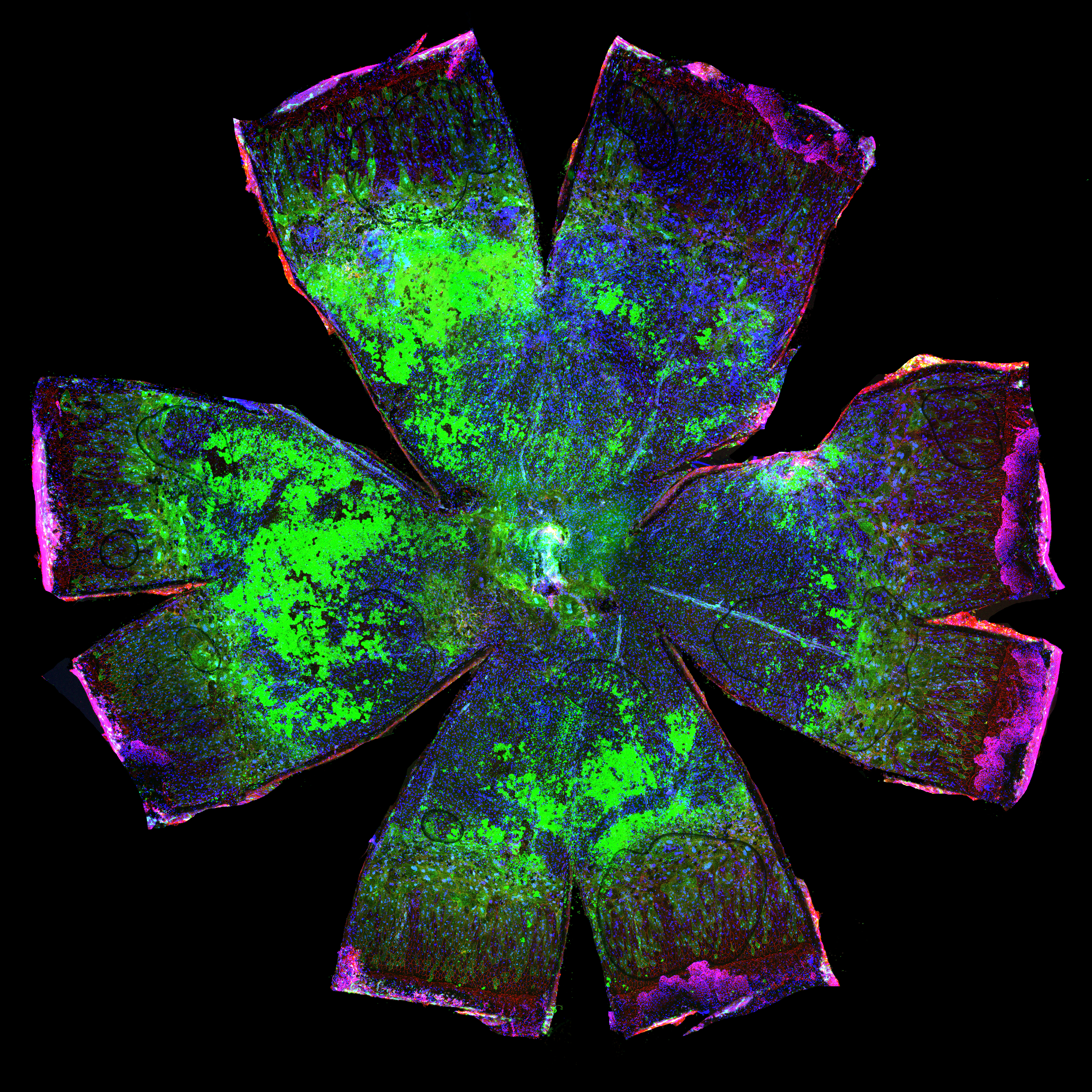

### Fig 4D: Full RPEFM-PBS-OD

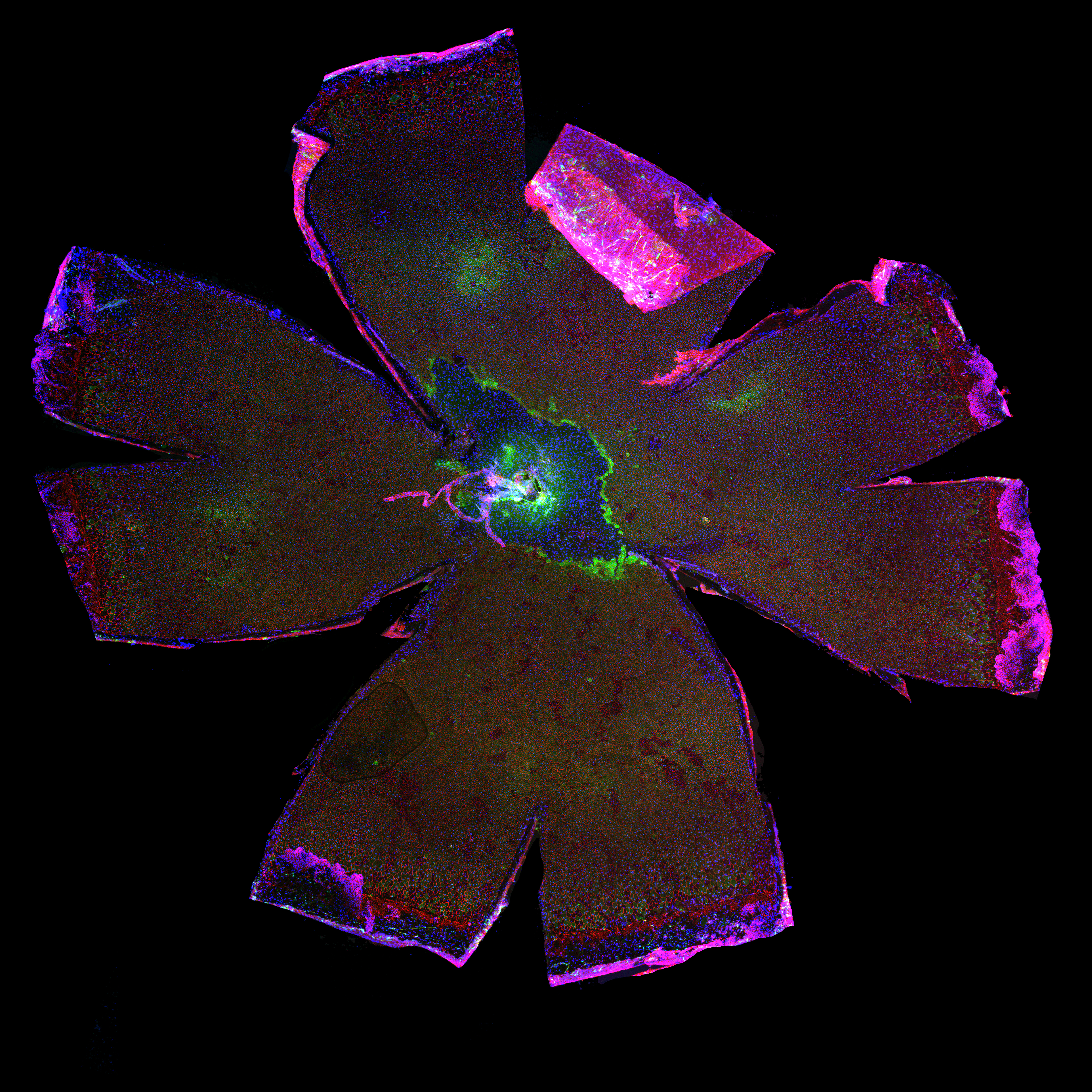

### Fig 4E: Full RPEFM-NaIO3-OD

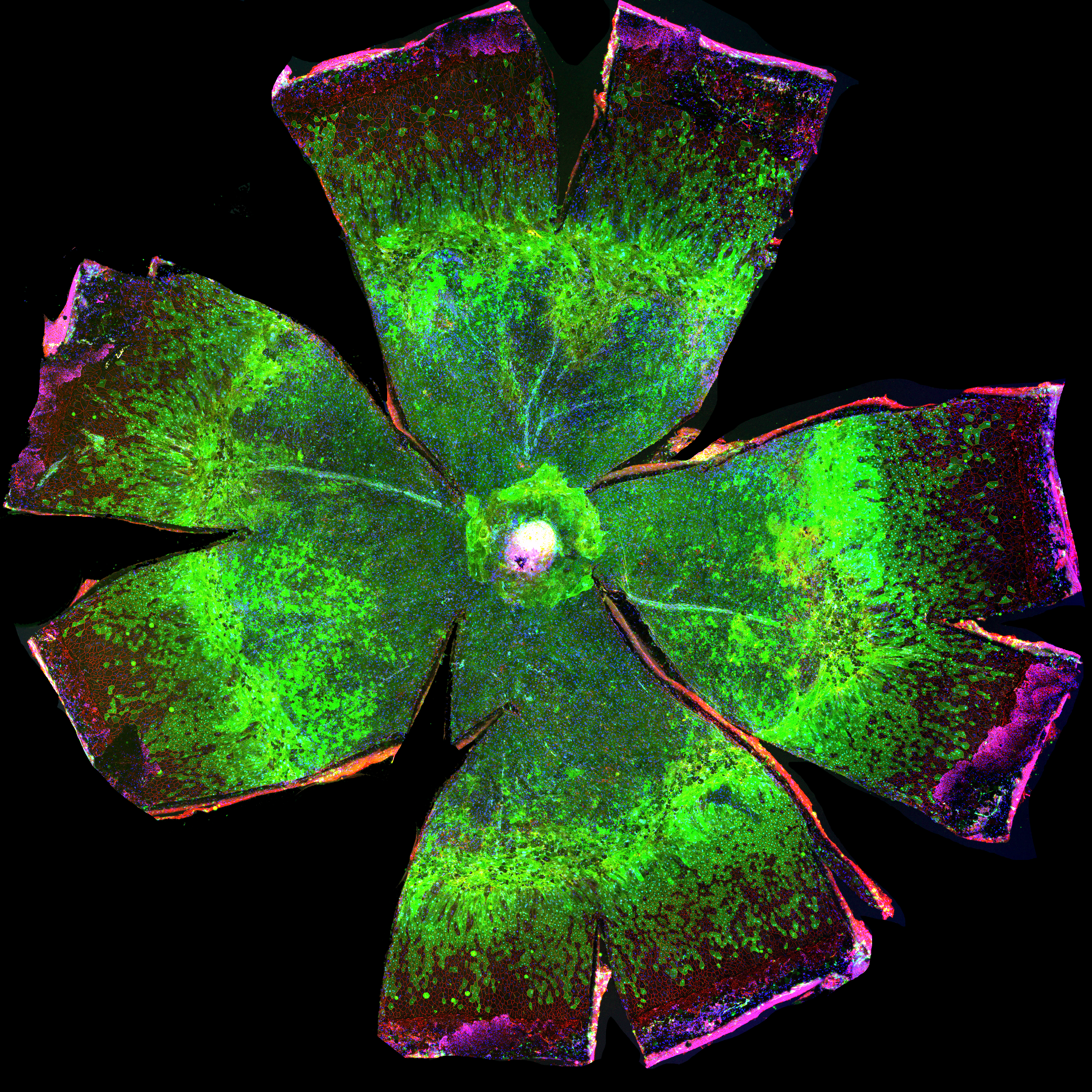

### Fig 4F: Full RPEFM-NaIO3-OD

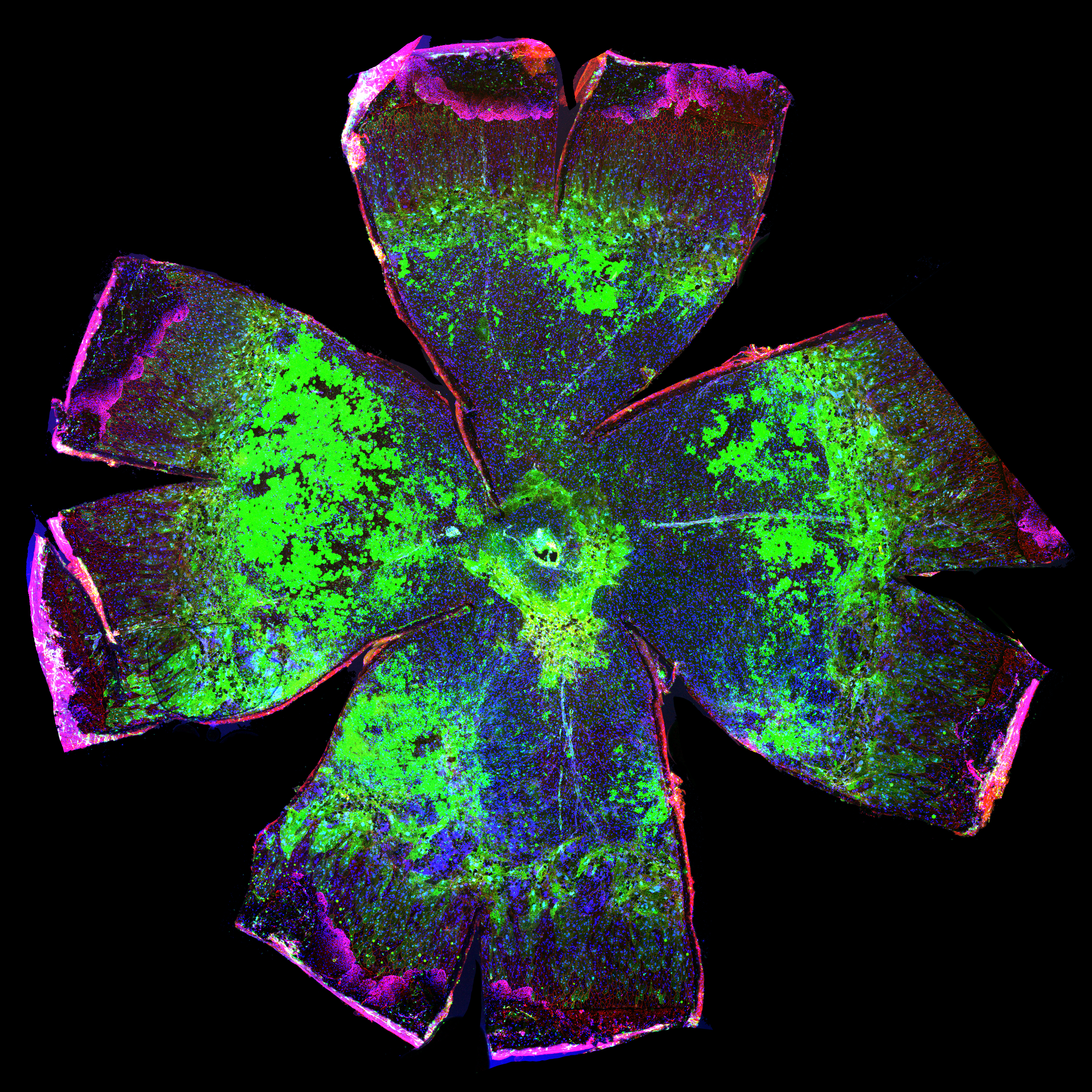
